# Comparative Evaluation of Targeted RNA Sequencing Protocols for Gene Expression Quantification With and Without Unique Molecular Indices (UMIs)

**DOI:** 10.1101/2025.01.27.635010

**Authors:** Annica Gosch, Cornelius Courts

## Abstract

Interest in forensic RNA analysis has increased over the last years. RNA molecules present in forensic samples can accurately be quantified via quantitative PCR (qPCR), however, due to the limited number of markers that can be assayed simultaneously per reaction, qPCR is less suitable for applications requiring gene expression quantification of large marker sets. Few years ago, massively parallel targeted RNA-sequencing (targRNAseq) allowing to simultaneously and accurately quantify several hundreds of markers has been added to the forensic genetic tool set. However, typical targRNAseq protocols include a multiplex-PCR-step to amplify selected targets which potentially introduces bias and limits accurate gene expression quantification.

Unique Molecular Indices (UMIs) have been invented to overcome this limitation and have been implemented in protocols from some vendors.

In this study, we compared two targeted RNAseq protocols assaying expression of a set of 121 forensically relevant mRNA biomarkers: The Ion Ampliseq targeted RNA sequencing panel (Thermo Fisher Scientific), which employs a multiplex-PCR without the use of UMIs, and the QIAseq targeted RNA panel (QIAGEN), which uses UMIs prior to multiplex amplification.

Both protocols were tested on replicated samples and dilution series and compared with respect to sensitivity and accuracy of gene expression quantification.

The UMI-based protocol exhibited decreased sensitivity in comparison to the non-UMI-based alternative, however, making use of UMI technology greatly improved gene expression quantification accuracy. We thus recommend the use of UMI-based protocols for targeted RNA sequencing for applications requiring accurate gene expression quantification.

## 1. Introduction

Analyses of differential gene expression to identify body fluids and/or organ tissues present in biological stains are routinely performed by several forensic laboratories worldwide and other potential applications of forensic RNA analysis are currently being explored.

Gene expression can accurately be quantified via quantitative PCR (qPCR) however, this method is less suitable for applications requiring the simultaneous analysis of large marker sets. Few years ago, massively parallel targeted RNA sequencing (targRNAseq) enabling simultaneous quantification of several hundreds of markers has been adopted for forensic molecular analyses. However, typical targRNAseq protocols include a multiplex-PCR to amplify selected targets which potentially introduces bias and limits accurate gene expression quantification. Unique Molecular Indices (UMIs), barcodes tagged to individual target molecules prior to PCR amplification and thus allowing to differentiate whether sequencing reads have been generated from a single or several distinct original target molecules, have been devised to overcome this limitation and have since been implemented in workflows and kits from some vendors.

Here, we assess whether the utilization of UMIs provides an advantage in targRNAseq for gene expression quantification in forensic samples. To this purpose, we compared two targRNAseq protocols to quantify the expression of a set of 121 forensically relevant mRNA biomarkers: The Ion AmpliSeq Library preparation kit (Thermo Fisher Scientific), in which multiplex PCR is performed without the use of UMIs, and the QIAseq targeted RNA panel (QIAGEN), which introduces UMIs prior to multiplex amplification.

## 2. Material studied, methods, techniques

### 2.1 Samples

Blood samples were obtained by fingerprick from four volunteers (2 males, 2 females) at two different time points of the day (donors A and B) or at a single time point (donors C and D). Samples were stored at -80°C immediately (donors A and B) or after a drying period at room temperature for 2 days (donor D) or for 2 days and 16 days (donor C, two samples taken directly after each other, stored for different time periods). Total RNA was extracted using the mirVana miRNA isolation kit (Thermo Fisher Scientific) following the manufacturer”s instructions including a lysis step for 30 min at 56°C. Genomic DNA was removed from the RNA extracts using the TURBO DNA free kit (Thermo Fisher Scientific) according to the manufacturer”s protocol. RNA concentrations were then quantified using the Qubit High Sensitivity Assay on a Qubit 2.0 Instrument (both Thermo Fisher Scientific).

### 2.2 Library Preparation and Sequencing

Sequencing libraries were prepared using the Precision ID Library kit with Precision ID IonCode™ Barcode Adapters (Thermo Fisher Scientific) and the QIAseq targeted RNA panel with the QIAseq Targeted RNA 96-Index HT L for Ion Torrent (QIAGEN) according to manufacturer”s instructions (“Ion AmpliSeq RNA libraries” protocol from the Ion AmpliSeq Library Kit Plus User Guide and “Ultra-low input and FFPE Sample v2” protocol, respectively). Custom primer panels targeting 121 forensically relevant mRNA markers for body fluid identification [1] and time of day estimation [2] were designed by and obtained from each manufacturer separately. For comparison of gene expression accuracy, sequencing libraries were prepared in triplicates (for two samples from donor A) or duplicates (for two samples from donor B) with an input amount of 20 ng total RNA per library.

For assessing the assay sensitivity, sequencing libraries were prepared in a three-point dilution series with input amounts of 20 ng, 5 ng and 0.5 ng (for three samples from donors C and D).

Libraries were quantified by qPCR (Ion Library TaqMan™ Quantitation Kit (Thermo Fisher Scientific) and QIAseq™ Library Quant Assay Kit (QIAGEN), respectively) and equimolarly pooled for sequencing on two Ion 530 sequencing chips. Templating and sequencing were performed using the Ion S5™ Precision ID Chef & Sequencing Kit on an Ion Chef and an Ion GeneStudio S5 system instrument, respectively (all Thermo Fisher Scientific).

### 2.3 Data analysis

Raw sequencing data analysis was performed on the Ion Torrent Suite platform (Thermo Fisher Scientific). For the Ion AmpliSeq-libraries, raw read counts were obtained as end-to-end-reads from sequencing data using the CoverageAnalysis plugin from the Ion Torrent Suite (Thermo Fisher Scientific). For the QIAseq libraries, raw read counts and raw UMI counts were obtained using the QIAseq targeted RNA panels gene expression pipeline on the web-based MyGeneGlobe platform (QIAGEN).

Raw read counts/raw UMI counts were rlog-transformed and Principal Component Analysis (PCA) was performed using the DEseq2 [3] package (version 1.42.1) in R (version 4.3.2).

## 3. Results

### 3.1 Accuracy

Replicate samples analyzed with the UMI-based protocol exhibited lower coefficients of variation than replicate samples from the protocol without UMIs (Figure 1a). Accordingly, in the PCA plot a tighter clustering of replicate samples and a better separation of distinct samples can be observed for the protocol with UMIs (Figure 1b).

**Figure 1:**
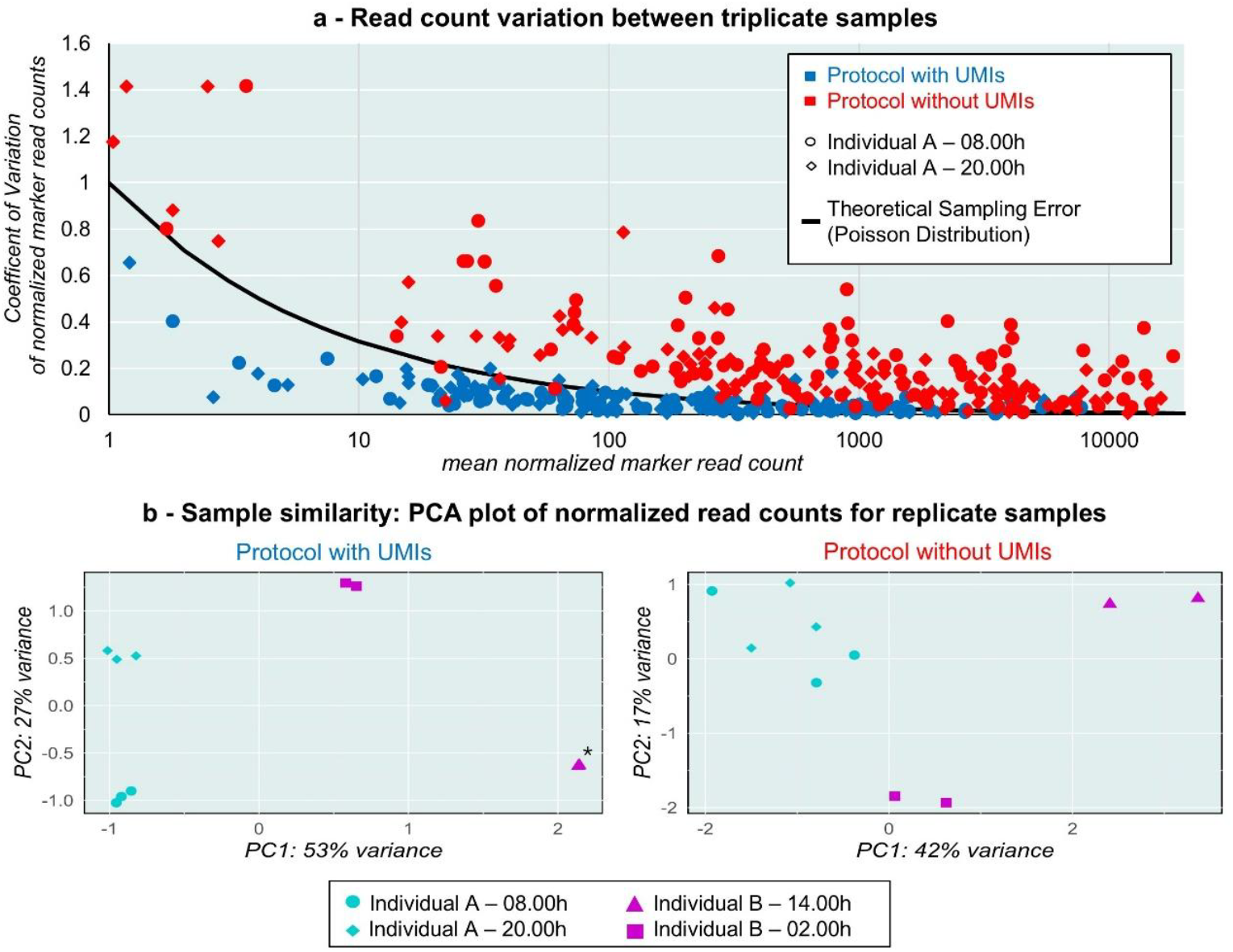
Comparison of gene expression quantification accuracy between sequencing libraries prepared with and without UMIs. **a** – Coefficient of variation between normalized marker read counts for two samples analyzed in triplicates with each protocol. **b** – Principal Component Analysis for samples from two individuals taken at two different time points each, analyzed in triplicates (donor A) or duplicates (donor B). ^*^Data points for individual B (14.00h) are directly above each other so that they cannot be visually separated.

### 3.2 Sensitivity

For the UMI-based protocol, marker detection was decreased with lower sample input amounts while this was not observed for the protocol without UMIs (Figure 2a). However, samples sequenced with the UMI-based protocol showed better separation according to biological properties even for the diluted samples, whereas samples from distinct donors were not separated in the PCA plot for the protocol without UMIs (Figure 2b).

**Figure 2:**
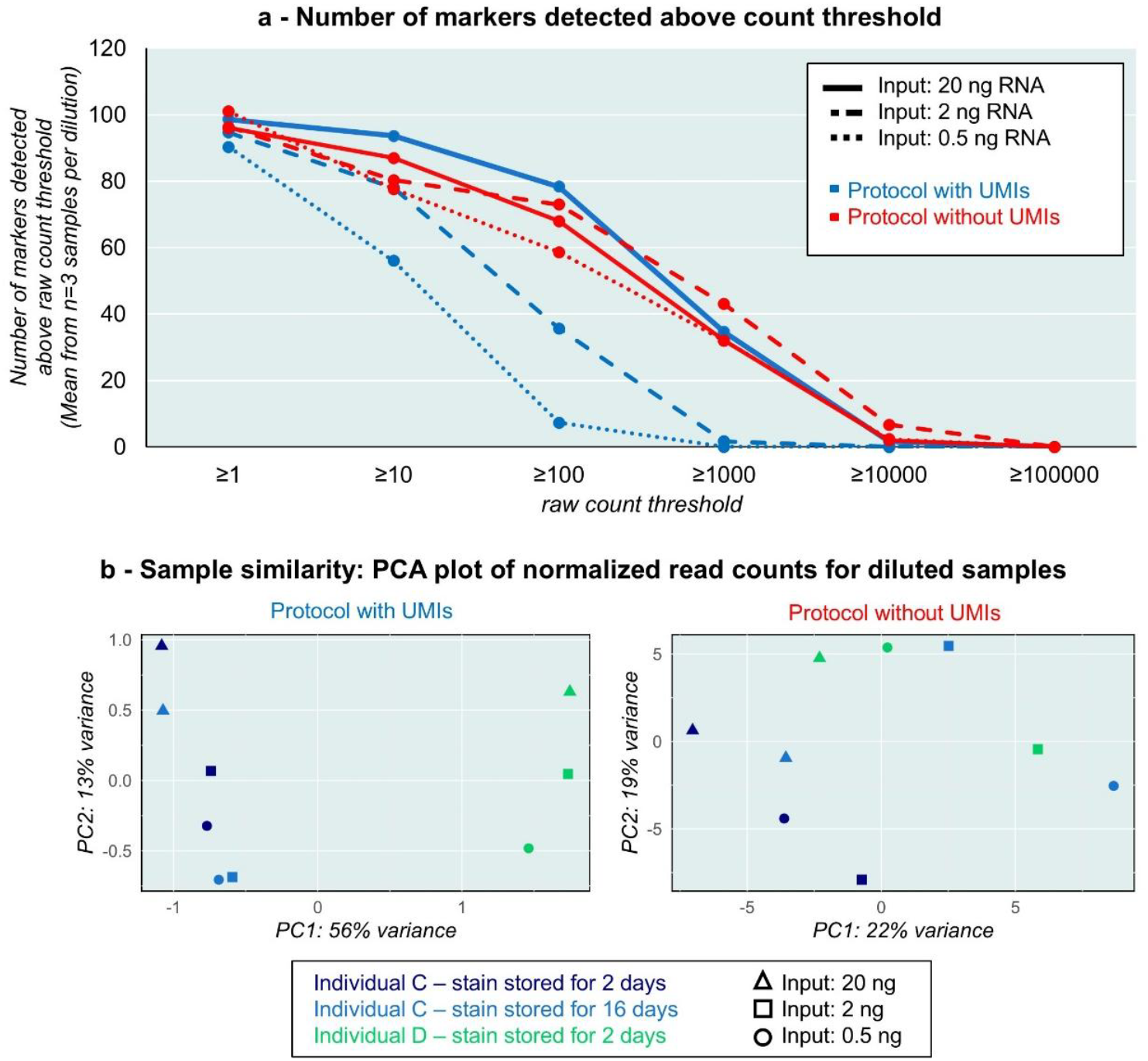
Comparison of gene expression quantification sensitivity between sequencing libraries prepared with and without UMIs. **a** – Number of markers detected above a raw read count threshold for sequencing libraries prepared from 20 ng, 5 ng and 0.5 ng total RNA with each protocol. Indicated values represent an average from three samples per dilution. **b** – Principal Component Analysis for samples from two individuals dried for two different time periods at room temperature (2 days and 16 days), analyzed in at RNA input amounts of 20 ng, 2 ng and 0.5 ng.

## 4. Discussion

The UMI-based protocol resulted in more accurate gene expression quantification compared to the library preparation protocol without UMIs, as evidenced by lower technical variation. This is in line with the theoretically expected advantage of UMIs reducing PCR-induced bias.

Marker detection sensitivity was reduced in the UMI-based protocol, which may be caused by loss of sample material in the molecular tagging step or during the bead clean-up steps in the library preparation protocol. However, while markers were still detected from very low input amounts with the protocol without UMIs, quantitative information from these measurements has to be considered less reliable as evidenced by a loss of the ability to distinguish distinct samples in PCA.

## 5. Conclusion

Replicate samples analyzed with the UMI-based protocol exhibited a lower variation than replicate samples analyzed with the protocol without UMIs. The detection sensitivity was reduced in the protocol with UMIs compared to the protocol without UMIs, however, biological differences (inter-individual differences, differences between sampling time points) were better captured with the UMI-based protocol. We thus recommend the use of UMI-based protocols for targeted RNA sequencing for applications requiring accurate gene expression quantification.

## 6. Acknowledgments

We are grateful to the anonymous sample donors for their participation in this study. We thank the Deutsche Forschungsgemeinschaft (DFG) for funding this study (CO-992/10-2).

